# Spontaneous onset of cellular markers of inflammation and genome instability during aging in the immune niche of the naturally short-lived turquoise killifish (*Nothobranchius furzeri*)

**DOI:** 10.1101/2023.02.06.527346

**Authors:** Gabriele Morabito, Handan Melike Dönertas, Luca Sperti, Jens Seidel, Aysan Poursadegh, Michael Poeschla, Dario Riccardo Valenzano

## Abstract

Turquoise killifish (*Nothobranchius furzeri*) are naturally short-lived vertebrates, recapitulating several aspects of human aging, including protein aggregation, telomere shortening, cellular senescence, and declined antibody diversity. The mechanistic causes of systemic aging in killifish are still poorly understood. Here we ask whether killifish undergo significant age-dependent changes in the main hematopoietic organ, which together could contribute to systemic aging. To characterize immune aging in killifish, we employed single-cell RNA sequencing, proteomics, cytometry and a functional in vitro assay. Our data indicate how old killifish display increased inflammatory markers, and while immune cells from adult killifish display increased markers of proliferation and replication-independent DNA repair in progenitor-like cell clusters, progenitors from old killifish display extensive markers of DNA double-strand breaks. In less than 10 weeks, killifish undergo several dramatic spontaneous aging-related changes in the immune niche, which could be functionally linked with its extensive systemic aging and serve as targets for anti-aging interventions.

## Introduction

Aging of the immune system has dramatic consequences on host physiology, reducing tissue regeneration and turnover^1^, lowering the establishment and maintenance of lasting immune memory^2^, increasing the risk for chronic infections^3^ and autoimmune diseases^4^. The causes of aging of the immune system are complex and still poorly understood. Both cell-intrinsic and cell-extrinsic factors have been proposed as contributors to aging of the immune compartment^5^. In mammals, aging of the immune compartment has been associated with increased DNA damage, to a shift in immune cell pools towards more myeloid over lymphoid cells (myeloid shift), as well as to an increased secretion of inflammatory mediators^6^. At the systemic level, aging of the immune system has been associated by a low-grade, chronic inflammatory state, called inflammaging^7^. Chronic inflammation, in turn, has been shown to exacerbate aging of the immune niche^8^.

The extent to which markers of immune aging displayed in mammals are shared in other vertebrates is to-date poorly understood^9^, with some evidence of immune senescence occurring in birds^10^.

By studying the cellular and molecular basis of immune aging in non-mammalian vertebrates, such as teleosts, we aim to uncover conserved mechanisms of immune aging and identify novel targets for interventions to enhance immune health during aging.

While studying immune aging in long-lived species could in principle help us discover the basis for extended homeostasis and immune health, naturally short-lived species offer the advantage of helping reveal the mechanistic basis for age-dependent homeostatic dysfunction.

African killifish of the genus *Nothobranchius* are characterized by a strikingly short lifespan for a vertebrate, with species living as short as 4 months both in captivity and in nature, after having reached sexual maturity in only 3-4 weeks^11,12^. Despite being short-lived, these fish display a wide range of age-dependent transformations after the onset of sexual maturation, including telomere shortening^13,14^, tumorigenesis^15^, neurodegeneration^16,17^, decreased capacity of tissue regeneration^18^, increased markers of cellular senescence in the heart^19^, loss of microbial richness in the intestine^20^, as well as decline in the diversity of the B cell antibody repertoire^21^. Given their wide range of conserved aging phenotypes, several killifish species – and especially turquoise killifish (*Nothobranchius furzeri*) – have been rapidly adopted in the past years as a powerful *in vivo* vertebrate experimental aging model system^22,23^. Whether organ-wide aging is caused by shared causes or whether different organs age independently from one another remains an open question, which is as relevant in killifish, as well as in other organisms, including humans^24^.

Here, we explored the changes in aging-dependent expression of canonical molecular markers of immune system aging in turquoise killifish, with a specific focus on the main hematopoietic organ. To gain insights into molecular changes specific to different cell types and functionally test immune responses in young vs. old killifish, we employ cytometry and a functional in vitro assay in immune cells stimulated with endotoxin (LPS). Furthermore, to characterize the age-dependent changes within the main hematopoietic organ and in the plasma of turquoise killifish, we combine different “omics” methods, generating a shared resource, including an R SHINY APP (KIAMO). Together, here we generate the first comprehensive reference dataset for immune aging in turquoise killifish and shed light on the mechanistic bases of spontaneous aging-dependent functional decline in a non-mammalian vertebrate.

## Results

### Plasma proteomics reveals increased markers of inflammation in aged killifish

Turquoise killifish evolved short lifespan, accompanied by organ-wide accumulation of aging markers^25^. Recent work focused on the B cell repertoire has shown that turquoise killifish display age-dependent decline in the antibody repertoire diversity, a bona-fide maker of aging of immune effector cells^21^. We asked whether turquoise killifish display systemic markers of immune aging by performing Tandem Mass Tag Mass Spectrometry (TMT-MS) in plasma from five adult (7 weeks old) and five aged-adult (16 weeks old) male turquoise (**Figure 1** and **Online Methods**). We favored males over females for technical reasons, due to their larger absolute size, which makes surgery more feasible (**Figure 1a**). We found that plasma from adult killifish has a distinct proteome from old killifish. To identify the functions associated with the plasma proteins that differentiate the most adult from old turquoise killifish, we performed Gene Ontology analysis, which showed proteolysis, coagulation, nucleosome structure and apoptosis as some of the main terms that increase with aging (**Figure 1**).

**Figure 1.**
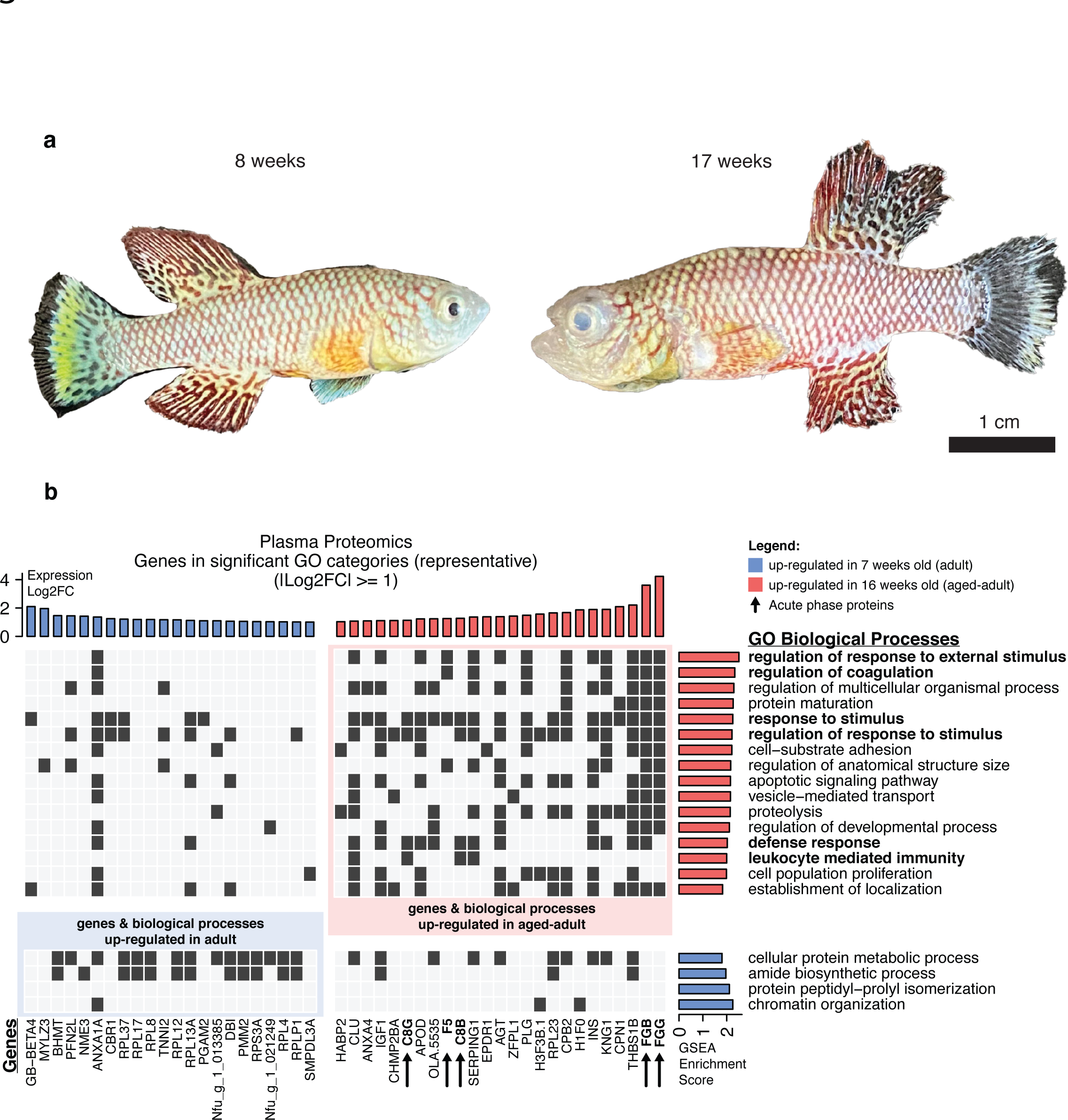
Plasma proteomics in young and old turquoise killifish show increased inflammatory terms in aged killifish. (A) Adult (8 weeks) and aged-adult (17 weeks) male turquoise killifish. B) Heat-map showing significantly enriched biological processes (GO categories) and all genes with |log2 fold change| ≥ 1 in the plasma of 5 young (7-week-old) and 5 aged (16-week-old) killifish. We considered BY corrected p-value ≤ 0.05 as a threshold for significance. Blue and red bars represent genes and GO categories respectively up-regulated in adult and aged fish. GO categories highlighted in bold play a role in inflammation. Dark pixels show GO category membership of the genes.

We then asked whether the aged killifish plasma shows enrichment for terms associated with immune system activation and inflammatory processes. None of the protein peptides identified by mass spectrometry are *bona fide* pro-inflammatory cytokines. However, we detected higher expression levels for several acute phase proteins (APPs) among the core enrichment genes of aged fish plasma, including complement proteins and coagulation factors, which are used as inflammatory markers in other teleost models^26,27^ (**Figure 1b**). In addition to the previous set of standard inflammatory markers, aged fish plasma shows also other core enrichment genes reported to be involved in aging and inflammatory processes: Carboxy-peptidase N1^28^, Annexin-A4^29^ and Apolipoprotein D^30^. Noteworthy, plasma from old fish showed higher levels of insulin^31^ and IGF1^32^ compared to non-aged adults, supporting an age-dependent systemic metabolic imbalance.

To further explore whether inflammation also affected cells residing in the main hematopoietic organ, we investigated the presence of histological markers of inflammation in the main hematopoietic organ, the teleost’s kidney marrow. We assessed the presence of tissue fibrosis, a marker of tissue inflammation^33^, performing a Fast green / Sirius red staining (**Online Methods**) in three 7-week-old (adult) and three 16-week-old (aged) killifish. We detected collagen deposition exclusively in the aged killifish tissues (**Extended Data Figure 1)**.

Together, plasma proteomics and histology revealed that killifish from the older age group display increased markers of inflammation both systemically, as well as in the main hematopoietic cell niche, in line with published results obtained in killifish livers, brains and intestine^20,34^.

### A single-cell atlas of the killifish main hematopoietic organ

To characterize the cellular composition of the killifish main hematopoietic organ (kidney marrow), we performed a single-cell RNA sequencing analysis using 10X technology. We sequenced cells from 6 adult male killifish. Five out of six samples showed high-quality reads and sufficient read numbers (**Online Methods** and **Extended Data Figure 2**). Therefore, we excluded one sample for the downstream analysis due to low read numbers. We performed reads clustering using the first 10 PCs of shared nearest neighbor graph construction, assessing a 0.3 resolution which provides the last most stable clustering (**Extended Data Figure 3 and Online Methods**). We annotated each cell cluster using differentially expressed, and cluster-specific, genes across the dataset (**Figure 2a** and **Online Methods**). The expression of cell-type specific markers confirmed that killifish have all the main vertebrate immune cell types, characteristic of both innate and adaptive immunity (**Figure 2b**).

**Figure 2.**
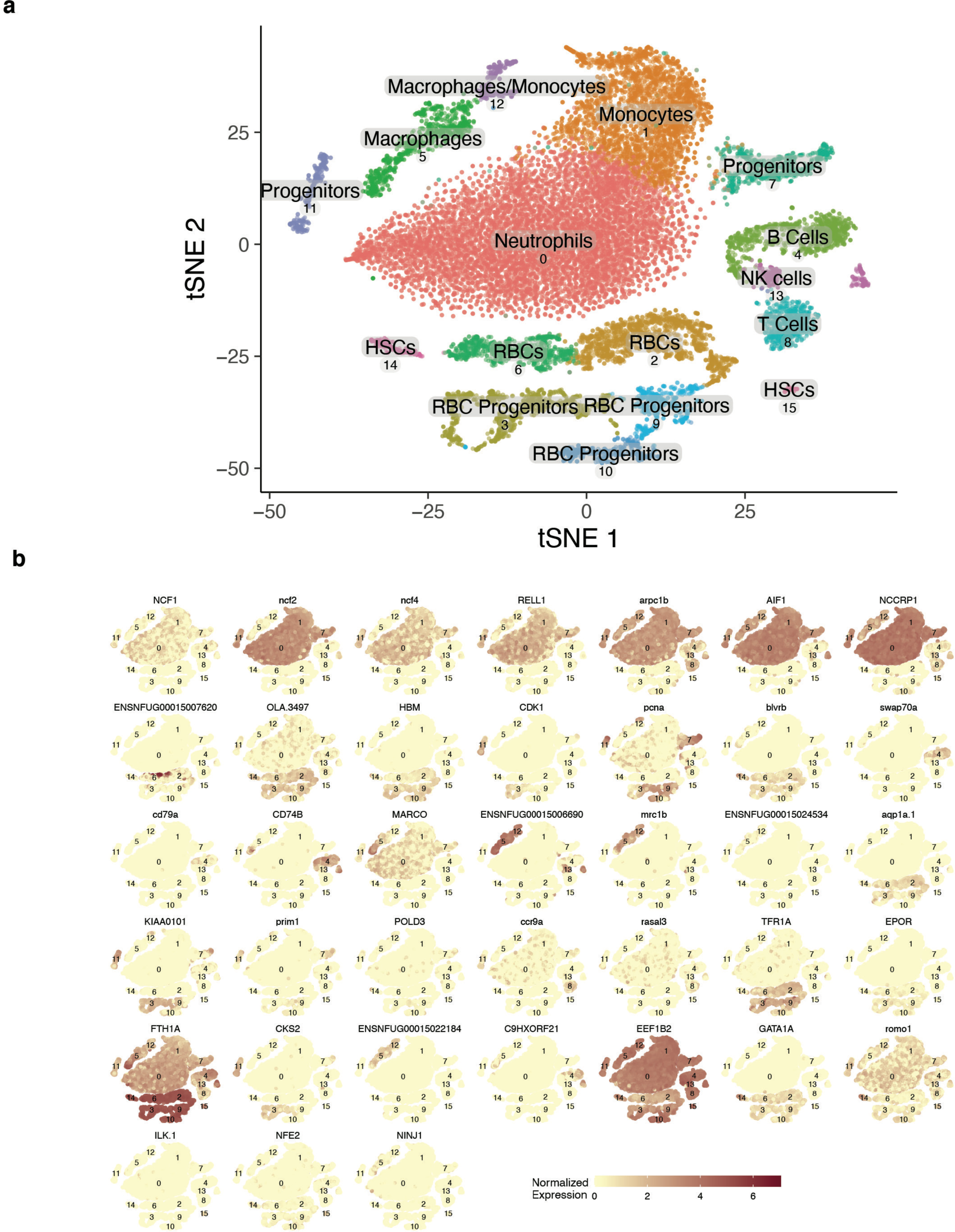
Cell composition in the main hematopoietic organ (kidney marrow) of turquoise killifish. a) tSNE plot of the cell clusters identified from single cell RNA Sequencing based on differentially expressed genes in 5 adult killifish (two 8-week-old, three 21-week-old). b) Selection of immune cell population-specific markers across cell clusters identified in the single cell RNA sequencing dataset.

### A cell cluster containing progenitors increases in relative cellularity within the kidney marrow during killifish aging

We asked whether cell composition in the kidney marrow varies as a function of age. However, in killifish there are no available antibodies that can be used as *bona fide* surface markers to distinguish specific immune cells clusters. Therefore, we took advantage of the variation in the biophysical properties among the cell populations extracted from the kidney marrow and used cytometry to identify separate cells based on cellular complexity (side-scatter) and cell area (forward scatter)^35,36^. Such cytometry-based approach allowed us to identify and count myelomonocytes-like, progenitors-like and lymphocytes-like cells across our samples (**Figure 3a** and **Figure 6a**). We extracted cells from the kidney marrow of 5 adults (8 week-old) and 4 aged-adults (17-week-old) killifish males and analyzed our samples using the Amnis ImageStreamX MkII Imaging Flow Cytometer. A cell cluster that typically contains progenitor cells within the fish kidney marrow appears to increase in relative abundance in aged fish (two-sided Wilcoxon ranked sum test p-value = 0.016) (**Figure 3b** and **Figure 6b**). However, the cell populations containing myelomonocytes-like, as well as lymphocytes, did not show any significant age-dependent changes (two-sided Wilcoxon ranked sum test p-values = 0.90 and 0.29, respectively). Together, our cytometric analysis broadly indicates that a cell cluster that typically contains teleost progenitors within the kidney marrow undergoes age-dependent changes between adult and aged-adult killifish.

**Figure 3.**
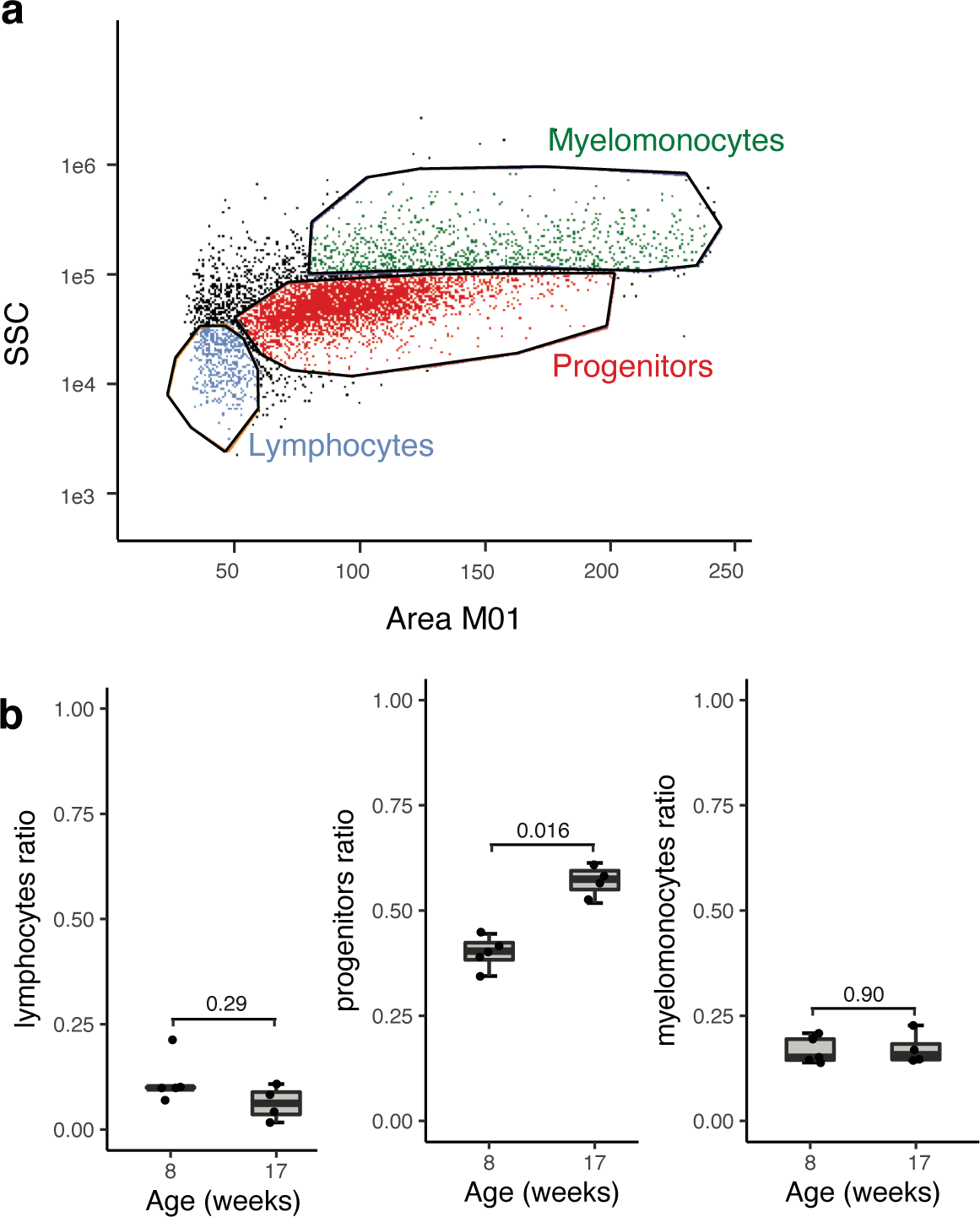
a) Dot plot of killifish kidney marrow single cell suspension obtained with Amnis ImageStreamX MkII Imaging Flow Cytometer and analysed in Amnis IDEAS v.6. SSC and Area M01 represent scatter properties and size of cells respectively. b) Boxplots obtained from the cytometric analysis showing cells proportions of myeloid, progenitors and lymphoid lineages across 5 adult (8-week-old) and 4 aged-adult (17-week-old) samples. All the statistics were done using non-parametric two-sided Wilcoxon ranked sum test.

### Proteomics in immune cells from the kidney marrow shows decreased proliferation and DNA repair during killifish aging

To assess which biological processes in the immune cells of the kidney marrow are affected the most during aging, we ran TMT-MS on proteins extracted from kidney marrow immune cells isolated from the same adult and old individuals used for plasma proteomics. Many more proteins were detected in head kidney proteomics than in plasma proteomics and several of them resulted differentially expressed during aging, showing a clear distinction between the two age groups (**Extended Data Figure 4**). We asked which biological processes differ the most with age in our dataset and we found a higher enrichment for proteins involved in cellular proliferation and “DNA repair” in adult individuals, while “cellular detoxification from toxic substances” and “negative regulation of chemotaxis” in aged-adult fish (**Figure 4a** and **Extended Data Figure 4**). Notably, in aged individuals we observed an enrichment for proteins involved in immune cell activation, such as: “interleukin-1b production” – a known pro-inflammatory cytokine^37^ –, “response to molecule of bacterial origin”, or “response to external biotic stimulus” (**Figure 4a-c** and **Extended Data Figure 4**). Hence, while components of DNA repair machinery in immune cells from aged individuals decrease compared to young-adult individuals, pro-inflammatory-related proteins are increased.

**Figure 4.**
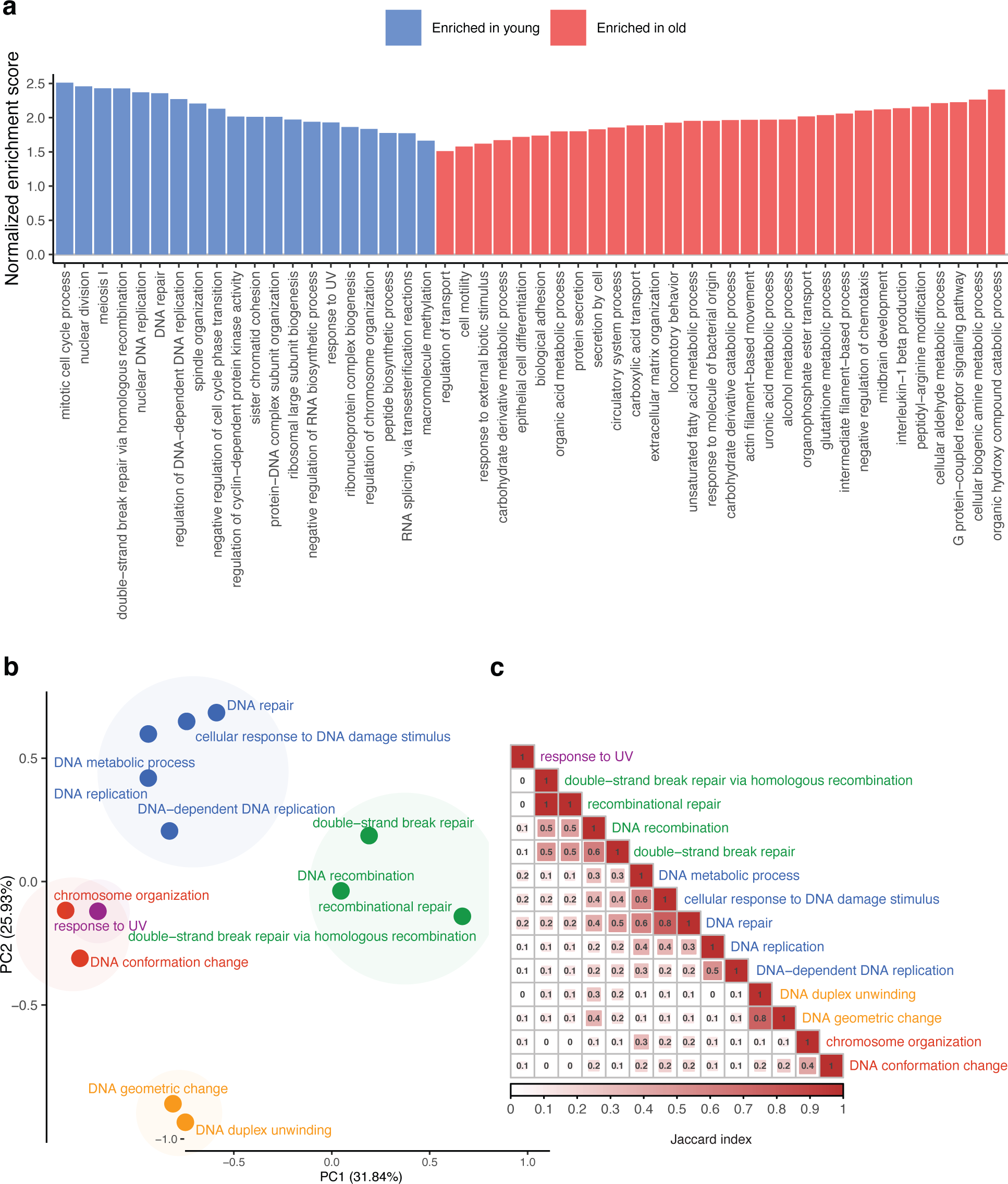
Aging-dependent changes in the proteome of the turquoise killifish hematopoietic niche. a) Barplot of the enriched biological processes in the immune cells isolated from the kidney marrow of 5 young (7-week-old) and 5 aged (16-week-old) turquoise killifish. We defined significance using BY-corrected p-value ≤ 0.05 as a threshold. Adults are in blue and aged-adults are in red. b) PCA plot of the GO processes represented by “DNA repair”, “response to UV” and “double strand breaks repair via homologous recombination” categories found enriched in young killifish and c) Correlation plot between all categories.

### Expression of DNA repair terms in young-adult killifish is specific to progenitors

To investigate which immune cell types were characterized by the expression of transcripts associated with the main GO categories enriched in our proteomics analysis between adult and aged-adult killifish, we focused on our scRNA Seq data. We found that the GO category “DNA repair” was associated with transcripts specific to all five progenitor cell clusters (**Figure 5**). We further found that the term “DNA replication” was not exclusive to the cell clusters showing markers of DNA repair. While progenitor clusters expressed both DNA replication and DNA repair proteins, we found additional cell clusters expressing DNA replication-associated genes, such as B and T cells, as well as clusters characterized as “hematopoietic stem cells” (HSCs). Together, our analysis suggests that DNA repair was not uniquely associated with DNA replication.

**Figure 5.**
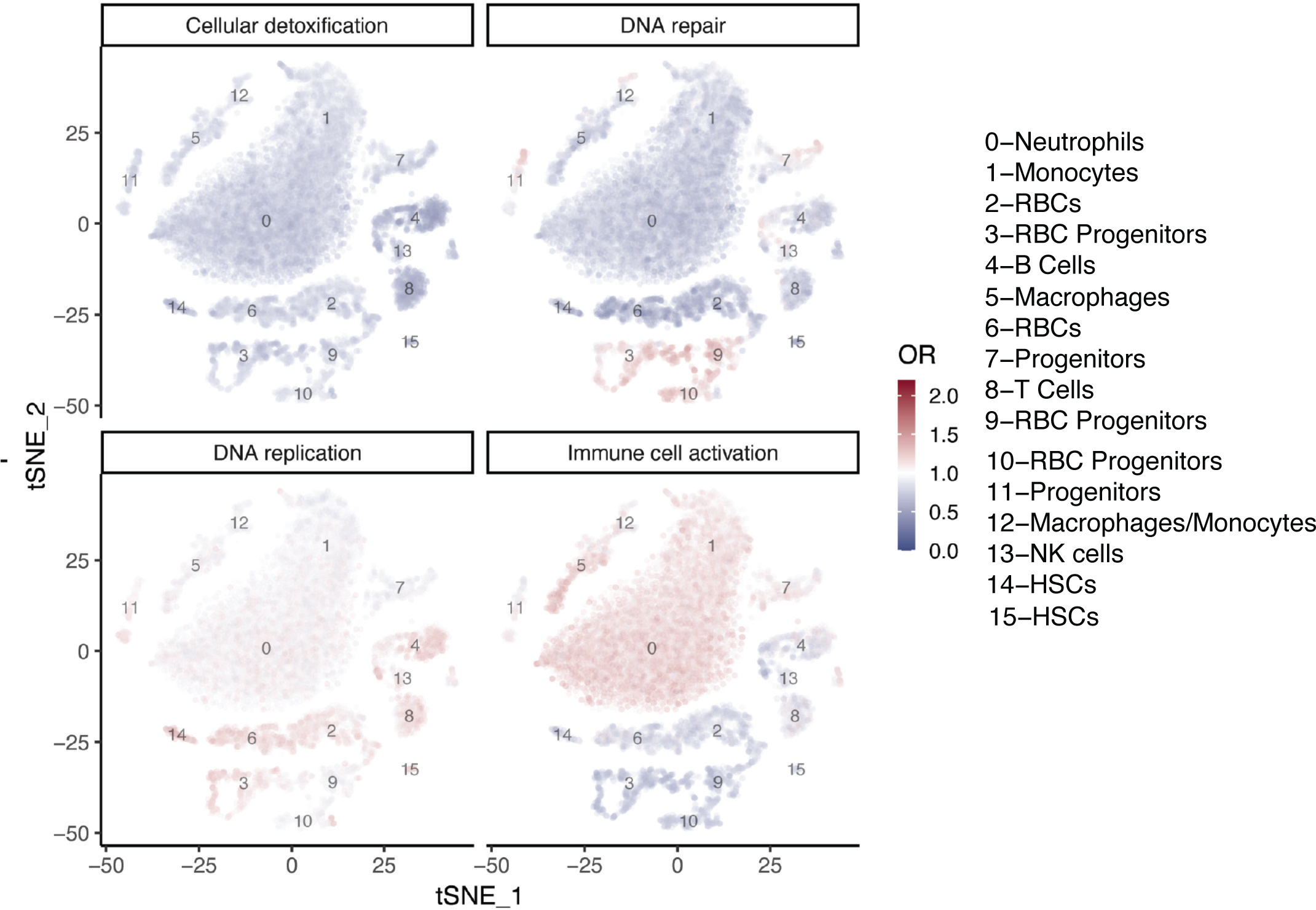
Cell-type specific expression profile of the enriched GO categories DNA repair and replication from proteomics data. a) tSNE plots showing enrichment of the genes belonging to ‘cellular detox’, dna repair, dna replication, and immune cell activation categories across cell-types. These four categories were chosen based on the results of proteomics dataset from the kidney marrow. Odds ratio (OR) is calculated by dividing the number of genes expressed in each category to the number expected by chance (see **Online Methods**).

### DNA damage increases in cells from the killifish kidney marrow during aging and doesn’t correlate with cellular proliferation

Since immune cells extracted from young-adult individuals, based on proteomics data, showed significant enrichment for cell cycle terms, we asked whether the higher expression of proteins associated with DNA repair in young-adult individuals was entirely explained by proliferation-dependent DNA damage. We setup an independent experiment using immune cells isolated from five 8-week old (adult) and four 17-week old (aged-adult) turquoise killifish. Since most of the DNA repair proteins over-expressed in young individuals participate in homologous recombination, a process used by the cells to specifically repair DNA double-strand breaks (DSBs), we used an antibody against γ-H2AX, which marks DSBs and overall genomic instability (**Online Methods**)^38^. We used the Amnis ImageStreamX MkII Imaging Flow Cytometer as a platform for a high throughput imaging of γ-H2AX signal^39^.

In each sample, to assess DSBs signal in DAPI positive cells, we used two different measures: the fraction of cells positive for DSBs over the total number of cells, calculated as the number of cells positive for γ-H2AX over the total number of DAPI positive-cells; and the median intensity of the fluorescent signal for γ-H2AX positive cells. Immune cells isolated from the kidney marrow of old fish contain more γ-H2AX positive cells and higher signal intensities compared to those derived from young fish, suggesting higher DNA damage signal in cells from old fish (**Extended Data Figure 8**) and confirming that in killifish, similar to other organisms, DNA damage increases as a function of age^40–42^.

Then, we asked whether cytometry confirmed the proteomics signal of higher cell proliferation in immune cells from younger fish. To this end, we used the DAPI signal of immune cells extracted from the kidney marrows of our aging cohorts^43^, which provides us with a measure of the phase of the cell cycle. Indeed, head kidneys from young fish had a higher fraction of cells in S/G2/M phase compared to head kidneys from old fish, supporting the higher proliferation signal in immune cells isolated from young fish.

If the signal for DSBs is uniquely associated with the cell cycle phase, we would expect cells under proliferation to carry a higher signal for DSB. We therefore asked whether the higher signal of DNA damage that we observed in immune cells from old killifish was associated with a higher fraction of proliferating cells (cells in S/G2/M phase). We found that cells from aged killifish, while having a smaller fraction of dividing cells compared to cells from young killifish, had a stronger signal for DNA damage (γ-H2AX) (**Figure 6c**-**e**). We then reasoned that an increased DNA damage signal in cells from aged head kidneys could result as the consequence of a) fewer dividing cells presenting a higher signal for DNA damage and/or b) non-dividing cells displaying markers of DNA damage.

**Figure 6.**
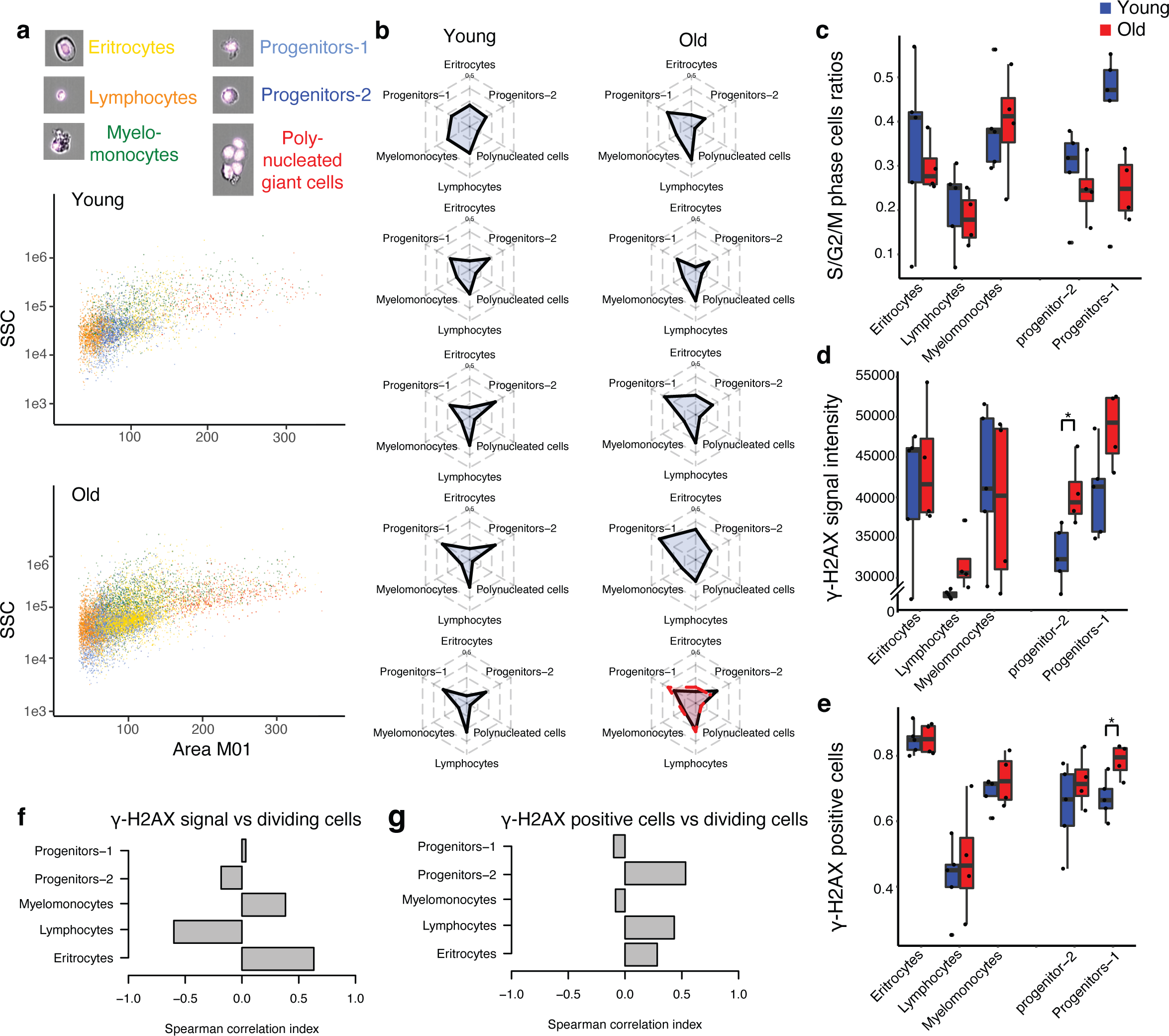
Machine learning analysis of *bona-fide* immune cells populations collected with ImageStream. a) Position of different *bona-fide* annotated immune cells populations in size scatter plots of adult and aged-adult fish. b) Cellularity changes in the hematopoietic niche of 5 adult and 4 aged-adult individuals. c) Dividing cells ratios across classified immune cells populations in the two aging cohorts d) y-H2AX signal intensities across classified immune cells populations in the two aging cohorts. e) y-H2Ax positive cells ratios across classified immune cells populations in the two aging cohorts. f-g) Correlation of y-H2AX measures with DAPI signal across classified immune cells populations in the two aging cohorts.

Since the DNA damage signal in replicating cells does not significantly change during aging (**Extended Data Figure 8c**), we consider unlikely that the increased age-dependent signal of DNA damage might be attributable to the pool of dividing cells. Our findings indicate that cell proliferation may not be the primary cause of DNA damage signals in cells from aged killifish (**Figure 6f-g**). Additionally, our results suggest that the heightened DNA damage signal and the reduced DNA repair signal in cells from older fish may not be linked to cell replication.

### Aging-specific γ-H2AX signal is specific to progenitors and does not co-localise with telomeres

Since we observed an age-dependent increase of DNA double-strand breaks, we asked which cell populations were mostly affected by this signal. Using a cytometry-based approach, we found that the cell cluster typically associated with progenitors showed most of the age-specific changes in γ-H2AX signal (**Figure 6d-e** and **Extended Data Figure 5**). To further explore whether the progenitor cells cluster carried the majority of the γ-H2AX signal, we adopted a machine-learning-based approach (**Online Methods**) to classify different cell types based on bright-field acquisitions provided via ImageStream. First, to automatically classify each cell type, we manually annotated all the main immune cell types based on their appearance as bright-field images. Then, we ran the model on a total of approximately 90K cells. To measure the concordance between manual classification and model prediction, we manually scored 19% of all cells from each cluster identified by the model (**Online Methods**), which gave us an accuracy value ranging from 75% (for Erythrocytes) to 96% (Polynucleated giant cells) (**Extended Data Figure 6**). Combining γ-H2AX antibody staining with our machine-learning based approach, we could support our finding that among all cell types, the progenitors cell cluster has the strongest signal for double-strand break within the killifish kidney marrow during aging. We further asked whether in the cells that – based on cytometry – we scored as progenitors, the signal for DSBs correlated with the ration of dividing cells. Supporting our previous finding, despite an increased signal for DSB in both progenitor cell clusters in old vs. young killifish, we did not observe an increased age-dependent ratio of dividing progenitors (**Extended Data Figure 5** and **Figure 6c**). Therefore, it appears that the accumulation of DNA damage during aging in the main hematopoietic organ in killifish is not associated with cell proliferation.

In our machine learning model, the progenitor population identified as progenitors II, shows in general less age-related differences in DSBs signal and dividing cells ratio compared to the progenitors I population. We acknowledge that this could be explained by the lower specificity of progenitors II classification in our model, which classifies around 50% of erythrocytes as progenitors II (false positives), hence buffering the signal in this population.

Next, we asked whether the aging-related increase of γ-H2AX signal is localized to telomeres, a typical source of signal for genomic double-strand break^44^. We performed an immunofluorescence assay on immune cells isolated from young and aged individuals using an antibody for γ-H2AX and a PNA probe for fluorescent in situ hybridization (FISH) against telomeric repeats (**Online Methods**). Noteworthy, we did not observe any clear localization of γ-H2AX in the telomeres across age groups, suggesting that the signal of DNA double-strand break in old killifish kidney marrows might not be specifically linked to telomere attrition (**Extended Data Figure 7**).

### Immune cells from aged killifish display an impaired functional response towards LPS stimulation associated with cellular senescence and are rescued by fisetin treatment

To test whether the molecular and cellular changes we see in killifish hematopoietic niche during aging are associated with an impaired immune cells functional response, we adopted an *in vitro* functional assay that measures immune cells activation in the presence of LPS, a potent immune stimulator^45^ (**Figure 7a**). We measured the response to LPS in immune cells extracted from young and old fish. LPS induced immune cell aggregates in a dose- and aging-dependent manner. At higher LPS doses, cluster numbers were lower in cells from the old killifish group (**Figure 7b**).

**Figure 7.**
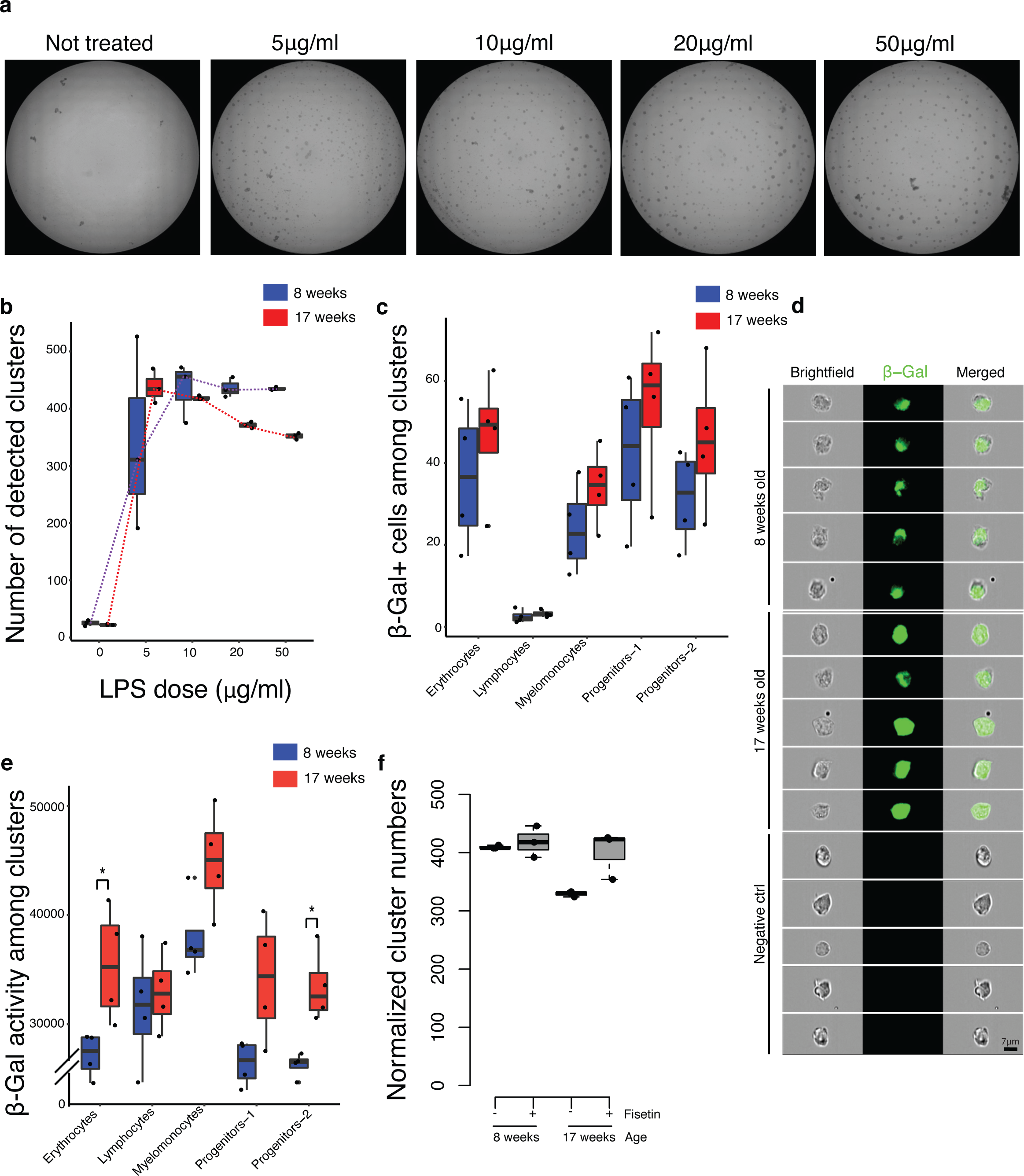
Machine learning-based analysis of *bona-fide* immune cell populations collected with ImageStream. a) Clusters formed by immune cells isolated from the hematopoietic niche of young and old fish incubated in the presence of different doses of LPS. b) Boxplot showing the effects of different LPS doses on the number of immune cells clusters formed (3 young vs. 3 aged fish). Normalized clusters number was calculated by subtracting the number of clusters obtained in the negative control from the number of clusters obtained after LPS stimulation. c-e) IF showing β-Galactosidase signal in immune cells isolated from the hematopoietic niche of 4 young and 4 old fish. β-Galactosidase positive cells ratios and signal intensities are shown by boxplots across classified immune cells populations in the two aging cohorts. f) Boxplot showing fisetin treatment effects on the number of immune cells clusters formed (3 young vs. 3 aged fish).

Since our previous results showed that cells isolated from old fish showed higher markers of DNA damage, we asked whether cells from old fish also presented typical markers of aging-dependent senescence. We performed a beta-Galactosidase assay and measured the signal using the ImageStream technology (**Figure 7c-e**). Similar to what already reported in mammals^46^, we observed an aging-dependent increase of senescence signal in cells from old killifish. Using our machine-learning-based classification, we could largely assign the age-dependent increase in senescence signal in the erythrocytes and progenitors clusters (**Figure 7e**). Next, we asked whether senescent signal in cells from old killifish contributed to the outcome of the LPS-induced cell cluster. To this end, we pre-treated for 24 hours immune cells isolated from young and old fish with the 15 μM of the senolytic fisetin^47^. Following fisetin treatment, we seeded the cells with 50 μg/ml of LPS. Remarkably, immune cells derived from old individuals that were treated with fisetin showed a number of clusters – upon induction with LPS – comparable to young cells; while immune cells from young-adult individuals treated with fisetin showed no difference compared to the control, untreated cells (**Figure 7f**). Together, these findings show that immune cells from old donors could return to a young-like behavior, in a cellular-based assay of immune induction, after pre-treatment with a strong senolytic.

### KIAMO: a platform to compare proteomics and single-cell RNA Seq data in turquoise killifish

To allow multi-omics comparison of plasma proteomics, head kidney proteomics and single-cell RNA Sequencing in the head kidney, we generated KIAMO (Killifish Immune Aging MultiOmics) https://genome.leibniz-fli.de/shiny/kiamo/KIAMO is a user-friendly shiny app that allows direct visualization across datasets in immune organs.

## Discussion

The immune system provides systemic surveillance, protecting organisms from a range of insults that come from the environment, e.g. tissue lesions, parasites and pathogens – as well as from internal processes, e.g. neoplastic transformations. Together, an effective immune system preserves homeostasis, while immune system malfunction is associated with a plethora of diseases, from chronic infections to cancer, neurodegenerative diseases, metabolic diseases, systemic inflammation, etc. During the aging process, the immune system becomes compromised as it is no longer able to effectively fight off pathogens and promote successful repair in damaged tissues. However, aging of the immune system is not solely associated with “lowered” immune effector function. Aging in the immune system is often characterized by hyperreactive immune responses^48^ and autoimmune dysfunctions, where the immune system is no longer able to properly discriminate self from non-self, resulting in increased auto-reactivity, ultimately leading to organ failure. Aging in the immune system is further characterized by a generalized, systemic, and chronic low-grade inflammatory state that has been named “inflammaging”^7^.

It has been recognized that aging of the immune system might be one of the main drivers of systemic aging-related functional decline at the organismal level across organisms^4,49^.

Cell-intrinsic mechanisms have been found to underlie aging-dependent failure in the hematopoietic stem cell niche^50^, including reduced regenerative capacity and age-dependent increase in genes involved in the myeloid vs. lymphoid lineage fate. Immune effector cells can also be impacted during aging, including lymphocytes^51,52^, as well as cells from the myeloid lineage^53,54^. However, the main molecular and cellular drivers of systemic aging of the immune system remain to date still largely unclear.

Our work aims to characterise the molecular and cellular signatures of vertebrate immune system aging. To this end, we adopt as a model system the African turquoise killifish (*Nothobranchius furzeri*), a naturally short-lived vertebrate with a lifespan ranging from four to eight months^11,23,25^, and a plethora of spontaneous age-dependent transformations, including neoplasias^55^, neurodegeneration^56^ and intestinal dysbiosis^20^. Recent studies have shown that aged killifish display markers of intestinal inflammation^20^ and undergo expansion of large B cell clones systemically, and a decrease in naïve B cells in mucosal organs^21^.

Here we explore whether systemic and immune aging in killifish is associated with molecular and cellular changes occurring systemically, as well as in the main immune niche. We found that killifish plasma shows age-dependent spontaneous increased abundance of insulin and IGF1, i.e. two major markers of aging-related metabolic and hormonal imbalance^57^. Furthermore, plasma from aged killifish is enriched with several markers of inflammation, including acute phase proteins. Strikingly, within a few weeks of life, turquoise killifish recapitulate conserved hallmarks of systemic aging associated with the onset of age-related diseases.

We further asked whether the kidney marrow in killifish undergoes significant age-dependent transformations. To this end, we first sought to chart the cell diversity within the main killifish hematopoietic organ, i.e., the kidney marrow, using single-cell RNA Sequencing. We found that killifish are characterized by the presence of all the main immune cell types found across vertebrates, including stem cells, several progenitor clusters, as well as differentiated cells from the erythroid, myeloid, and lymphoid lineages. Our cytometric analysis suggests an increased presence of progenitor cells during aging, coupled with decreased overall proliferation, supported by both cytometry and proteomics. Together, our results suggest the presence of an aging-dependent increased pool of progenitor cells, which could be compatible with impaired differentiation into effector cells.

However, given the current lack of reliable surface markers in killifish that might help distinguish among the various progenitor cell populations, we cannot yet address whether specific subclasses of progenitor cells (e.g. myeloid, erythroid or lymphoid) increase during aging more than others.

Our results further indicate that cells from the killifish kidney marrow present increased markers for double-strand DNA breaks and replication stress in aged vs. non-aged adults, while proteomics markers of active cell cycle and DNA repair are increased in cells from the non-aged adult killifish. Markers of DNA damage can occur in association with active cell replication. However, our results support that the increased signal for DNA damage in immune cells from aged individuals is overall independent from cell replication. Noteworthy, we found that the signal of DNA double strand break was not localized at telomeric regions. Furthermore, we could find cytometric support towards a specific increased DNA damage signal in progenitors from aged killifish.

Together, our results in killifish support that progenitors undergo DNA damage and possibly replication stress upon aging, while they might be protected by active DNA repair mechanisms in non-aged, young adults. Hence, contrary to the expectations that hematopoietic progenitors would be geno-protected, we found that HPCs from old killifish carry higher levels of DNA damage compared to all other cell type in the hematopoietic organ, consistently with what recently shown in aged mammalian hematopoietic stem cells^58,59^.

Remarkably, in about two months-time, i.e., from non-aged adults to aged-adult killifish, we observe the rapid onset of extensive cellular markers of immune aging, suggesting that the main killifish hematopoietic organ undergoes rapid and dramatic age-dependent changes. Finally, our findings show that targeting senescent cells extracted from old killifish restores immune responses to LPS, a powerful immune stimulant. While our preliminary in vitro results show a putative role for senescence in immune cell functional changes during killifish aging, whether the molecular and cellular changes occurring in the killifish kidney marrow during aging lead to systemic functional consequences remains still to be assessed.

The main hematopoietic organ of aged killifish presents markers of inflammation, reduced proliferation, reduced DNA repair, as well as increased non-telomeric DNA damage. Our work provides an extensive resource for aging omics in killifish, and we developed an R Shiny app (KIAMO) to access and visualize proteomics and single-cell transcriptomic data in in young and old killifish immune organs.

Given their experimental accessibility – e.g., via genome editing – and a rapidly expanding scientific community, killifish represent a powerful naturally short-lived vertebrate for *in vivo* studies of aging of the immune niche.

## Acknowledgments

The authors declare no conflicts of interests.

Mass spectrometry proteomics analysis was performed in the Proteomics facility of the Max Planck Institute for Biology of Ageing. Xinping Li advised on sample preparation, Ilian Atanassov performed initial computational proteomics analysis. Janine Altmüller from the Cologne Center on Genomics (CCG) conducted led the scRNA Sequencing. The Life Science Computing Core at the FLI hosted KIAMO and the Imaging facility of the FLI facilitated image acquisition. We thank all the members of the MPI-AGE and FLI fish facilities for their key contribution in running our fish cohorts.

We thank the Max Planck Institute for Biology of Aging, the Max Planck Society, the Leibniz Institute on Aging, the DFG Collaborative Research Center SFB1310, and EMBO postdoctoral fellowship (HMD) for the financial support for this project. We thank Fabrizio d’Adda di Fagagna’s group at IFOM for their key input on the FISH experiment. We thank all past and current members of the Valenzano lab for their continuous critical input and support on this project.

## Authors contribution

GM and DRV conceptualised the experiments. GM performed all histological experiments, provided the proteomics facility with all the tissues for proteomics analysis, ran the ImageStream experiments, and generated the figures. GM and LS ran the cluster forming assay. HMD analysed all the omics data, generated KIAMO and contributed to the figures. JS, AP and MP generated the scRNA Seq dataset. DRV conceived the project and wrote the manuscript with the help of GM and HMD. All authors provided intellectual contribution to this project.

---

**Extended Data Figure 1.**
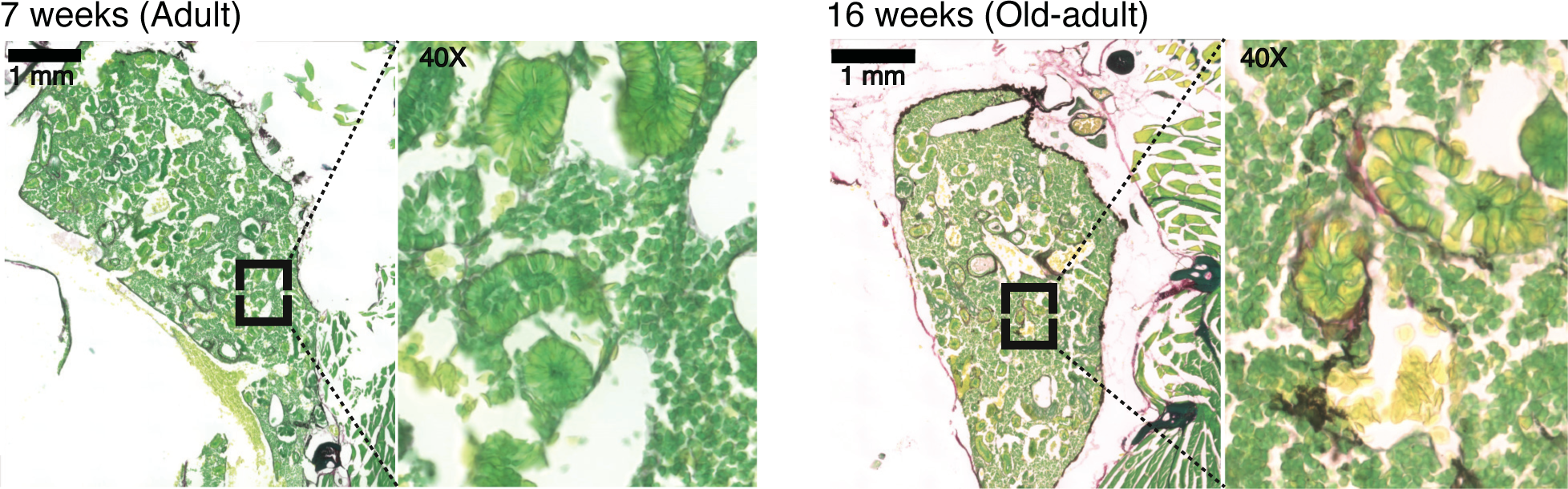
Kidney marrow displays aging-dependent fibrosis signatures. a) Fast green/ Sirius red staining of 3 adult and 3 aged-adult killifish kidney marrows. All the images show in red collagen (firbrotic area) and in green the rest.

**Extended Data Figure 2.**
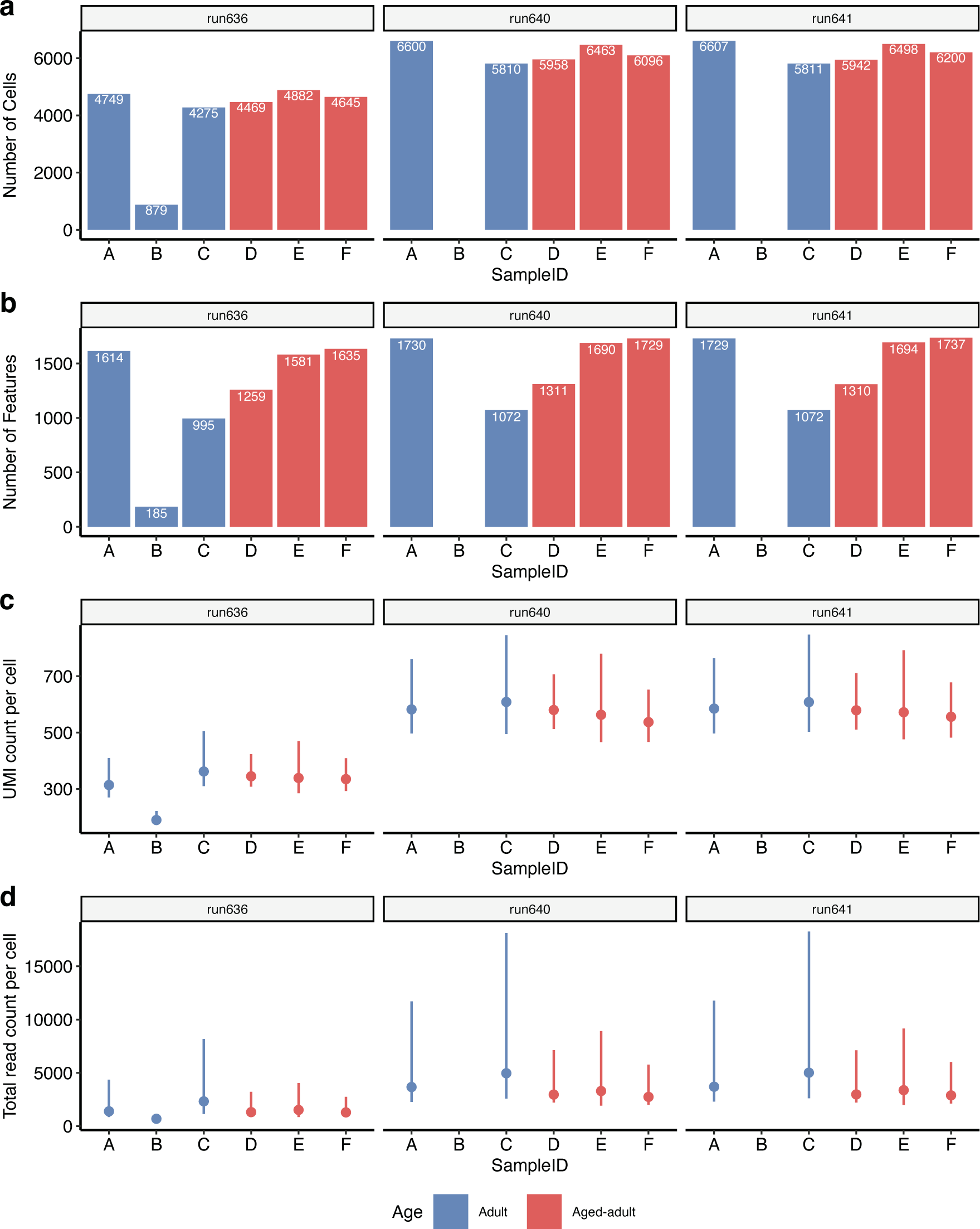
Analysis of reads quality across single cell RNA sequencing samples.

**Extended Data Figure 3.**
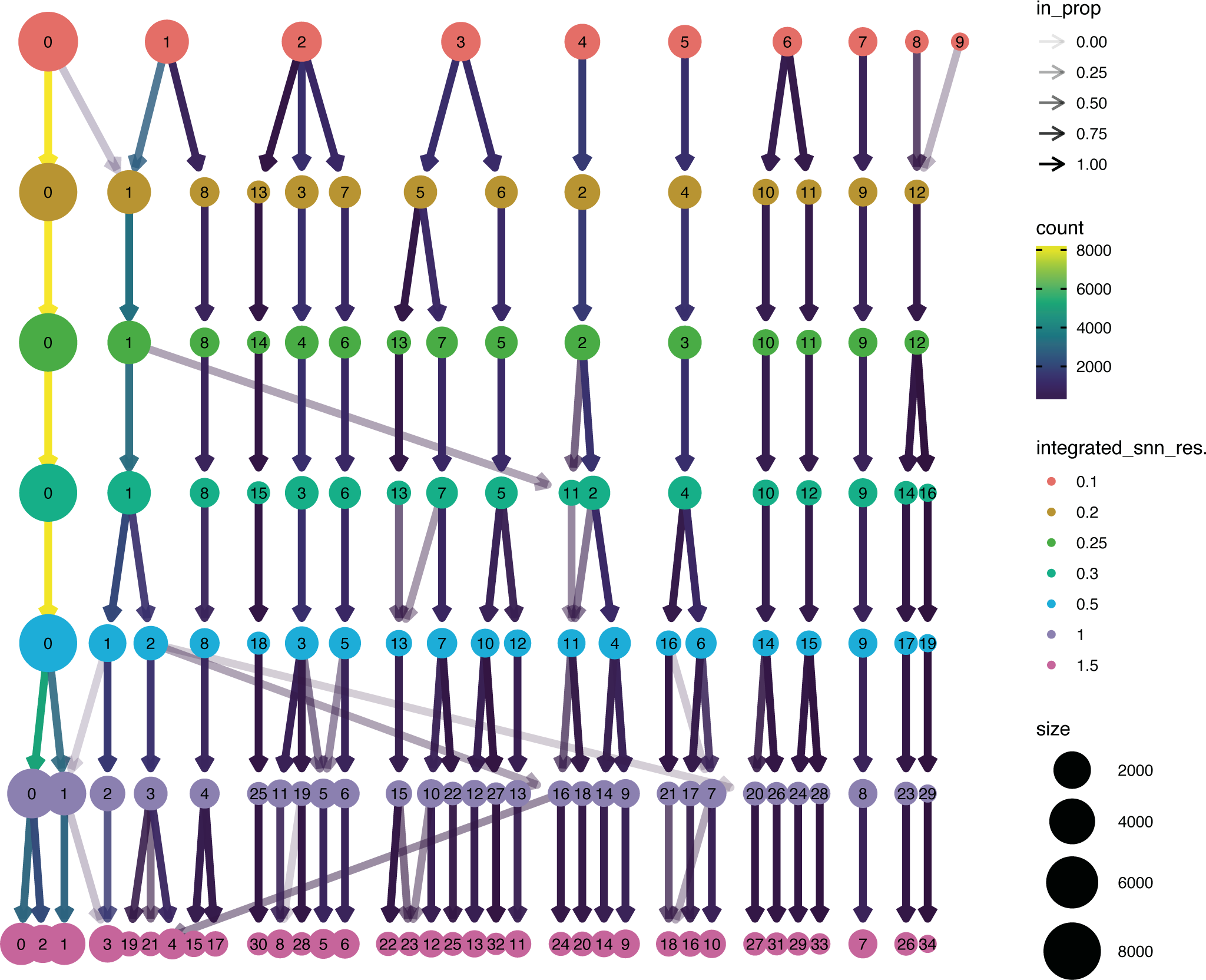
Clusters identified at different resolutions in single-cell RNA sequencing.

**Extended Data Figure 4.**
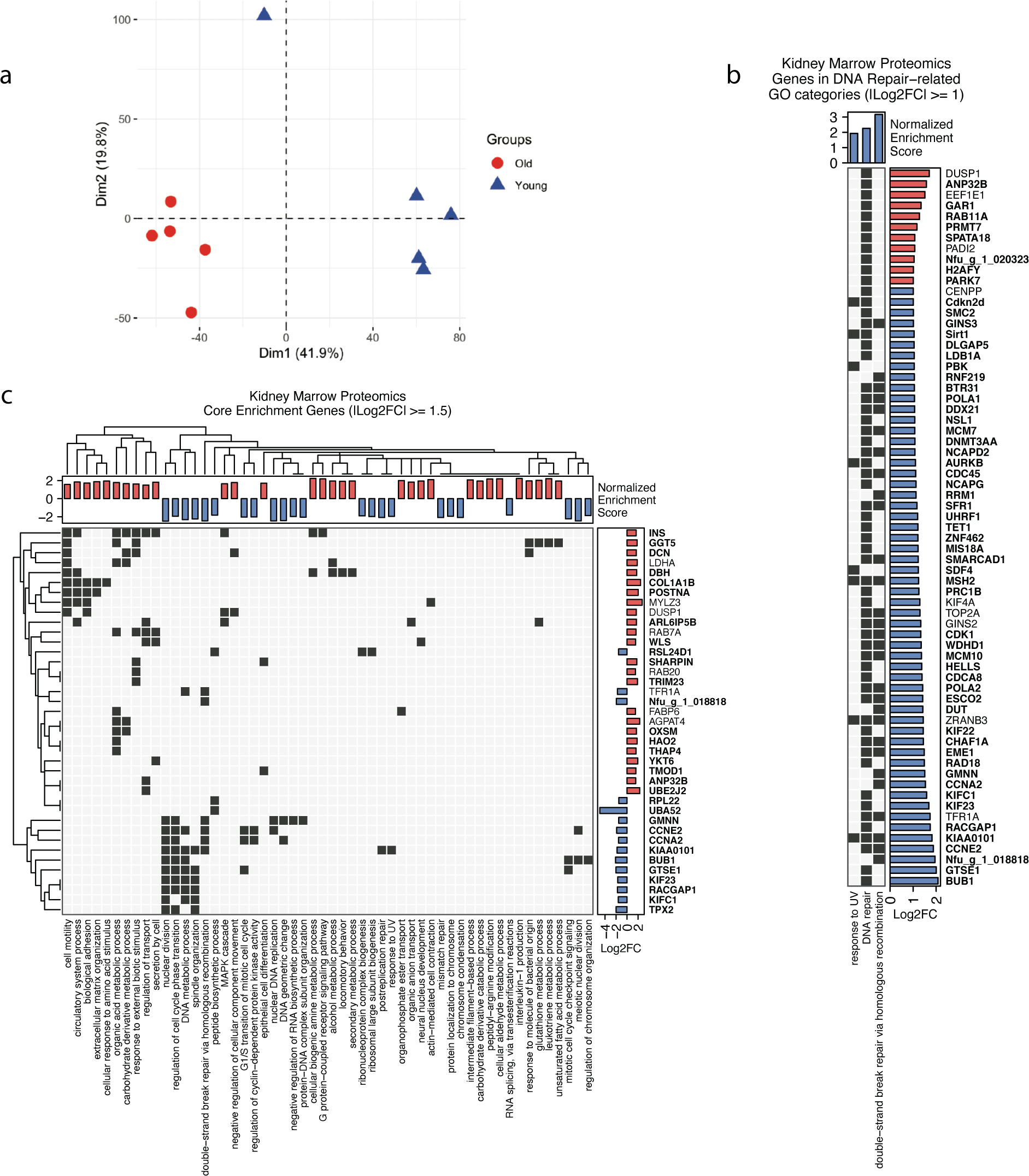
Significant biological processes and DNA repair proteins characterizing aging in the primary immune niche in turquoise killifish A) PCA plot, using all kidney marrow proteins in VSN-normalized, log2 transformed and scaled expression data, showing clustering of adult and aged-adult samples. B) Heat-map showing kidney marrow proteomics enriched DNA repair associated GO categories (“DNA repair”, “response to UV” and “double strand break repair via homologous recombination”) and their representative genes with a |log2 fold change| ≥ 1 in the kidney marrow of 5 adult (7 week old) and 5 aged (16 week old) killifish. We considered BY corrected p-value ≤ 0.05 as a threshold for significance. Blue and red bars represent genes and GO categories respectively up-regulated in adult and aged fish. GO categories highlighted in bold are significant. C) Heat-map showing significantly enriched biological processes (GO categories) and all genes with |log2 fold change| ≥ 1 in the kidney marrow of 5 adult (7 week old) and 5 aged (16 week old) killifish. We considered BY corrected p-value ≤ 0.05 as a threshold for significance. Blue and red bars represent genes and GO categories respectively up-regulated in adult and aged fish. GO categories highlighted in bold are significant. Dark pixels show GO category membership of the genes.

**Extended Data Figure 5.**
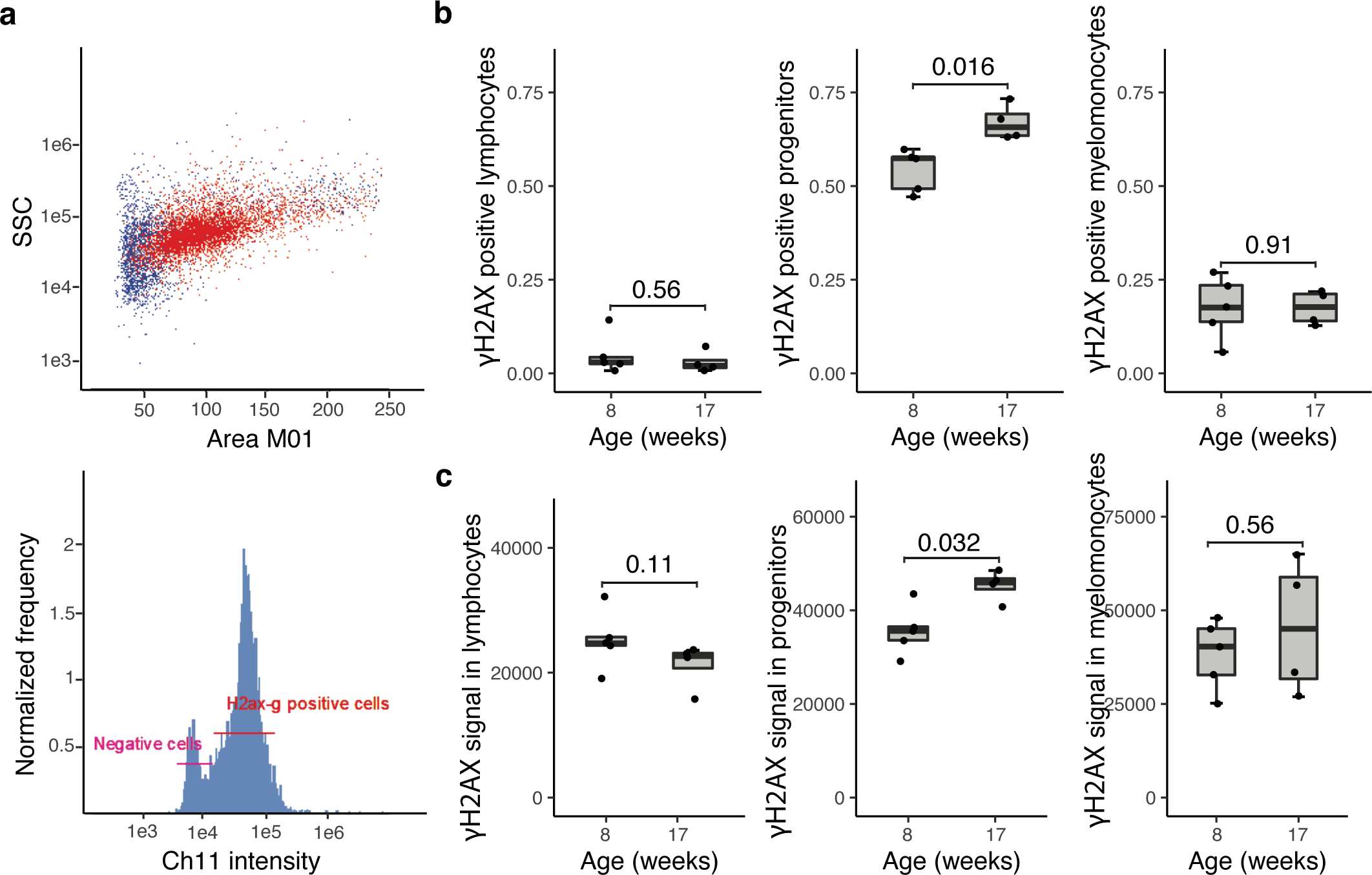
Cytometry measure of γH2AX signal in different immune cells clusters a) Above is shown the dot plot showing the overlap of whole single cells suspension (blue) and γH2AX positive cells (red) within killifish kidney marrow immune cells. Below is shown the histogram of typical γH2AX positive and negative cells frequencies pattern within across the samples analysed. b-c) Boxplots obtained from the cytometric analysis showing γH2AX signal deconvolution in myeloid, progenitors and lymphoid lineages across 5 adult (8 week old) and 4 aged-adult (17 week old) samples. All the statistics were done using non-parametric Wilcoxon rank sum test (two-sided).

**Extended Data Figure 6.**
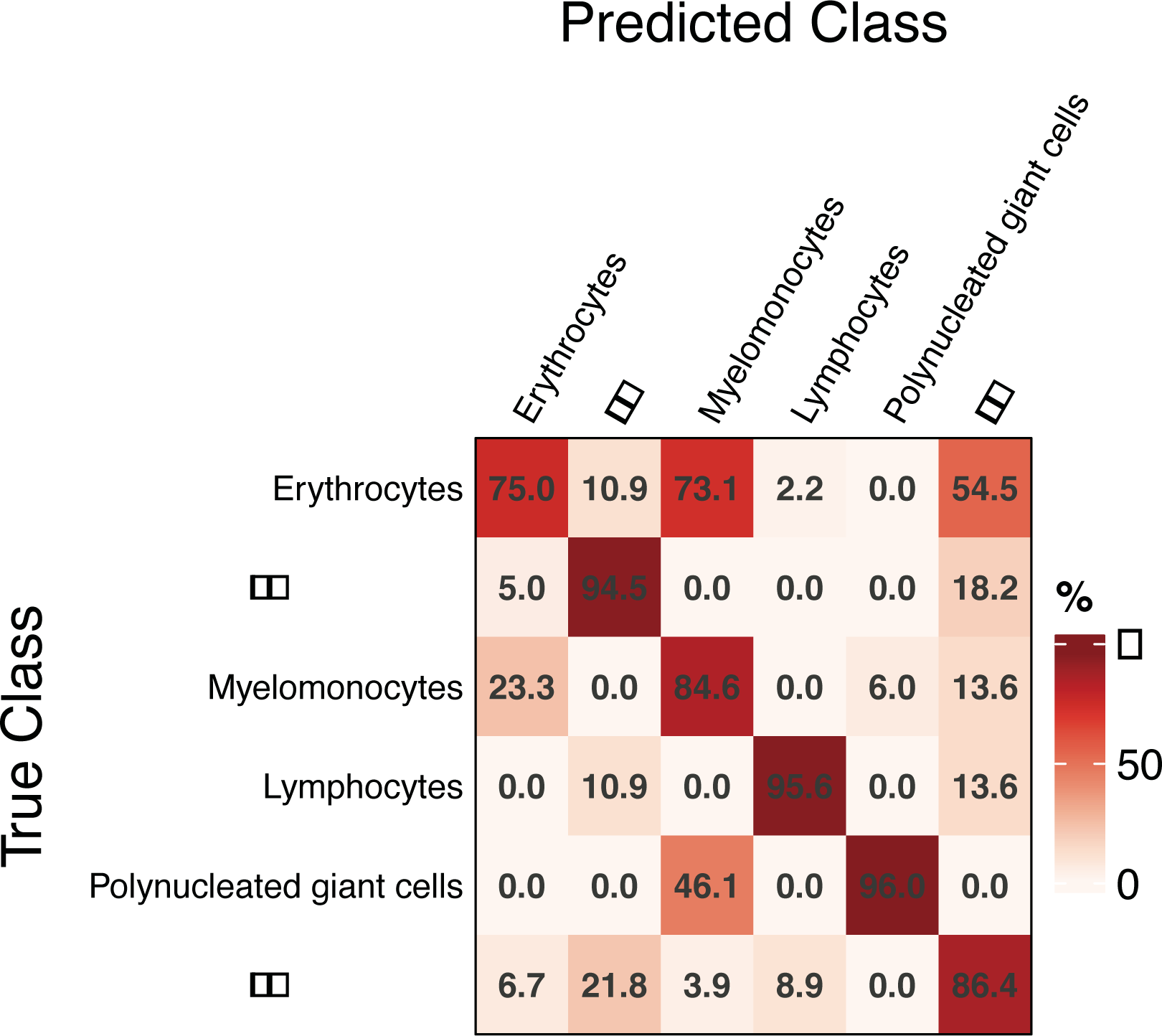
Accuracy matrix of the predicted immune cells clusters fit within true classified immune cells population.

**Extended Data Figure 7.**
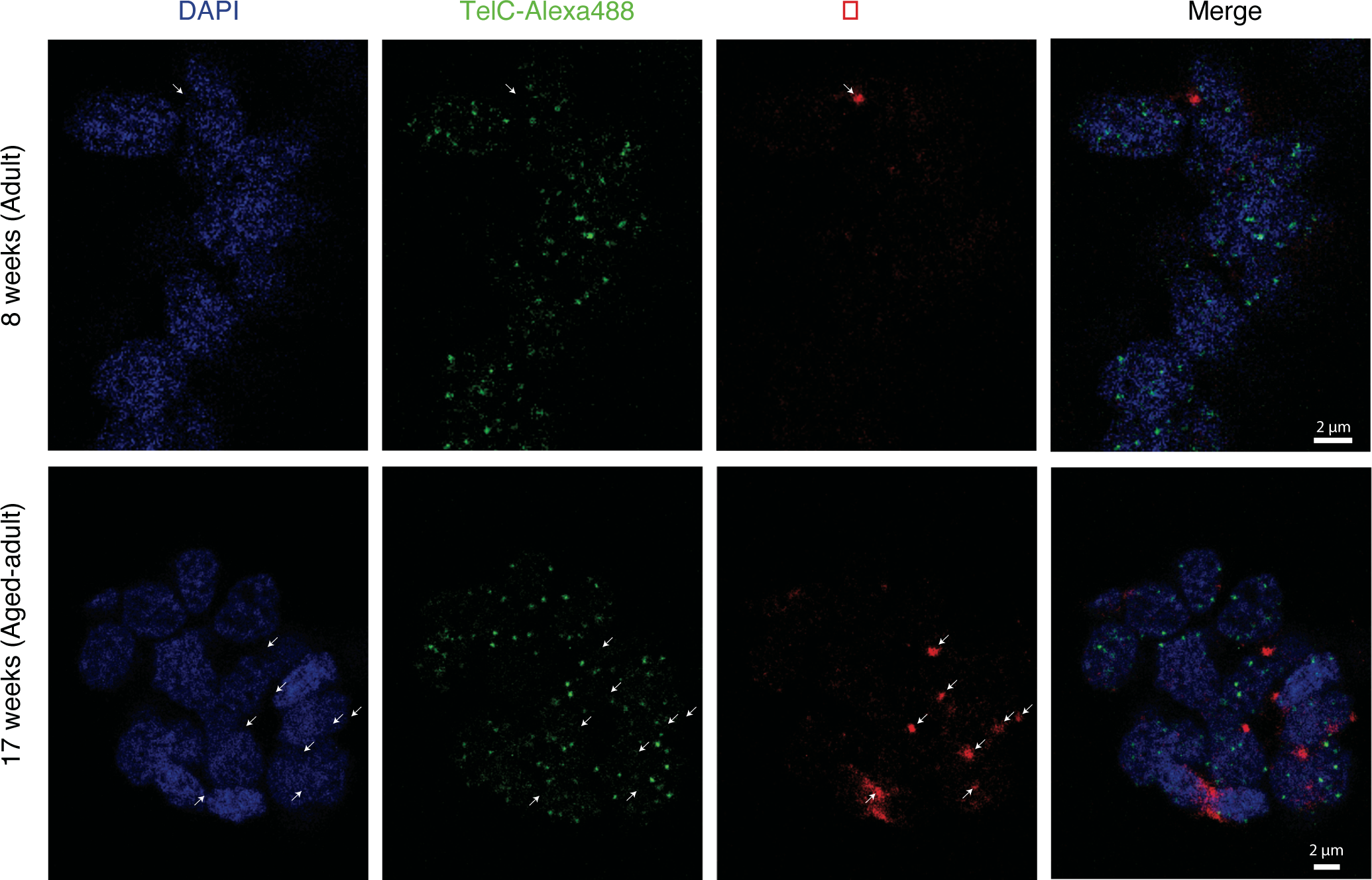
Co-staining of γH2AX and telomeric probes. a) IF/FISH showing localization of γH2AX and Telomeres within immune cells isolated from young-adult (8 week old) and aged-adult (17 week old) individuals.

**Extended Data Figure 8.**
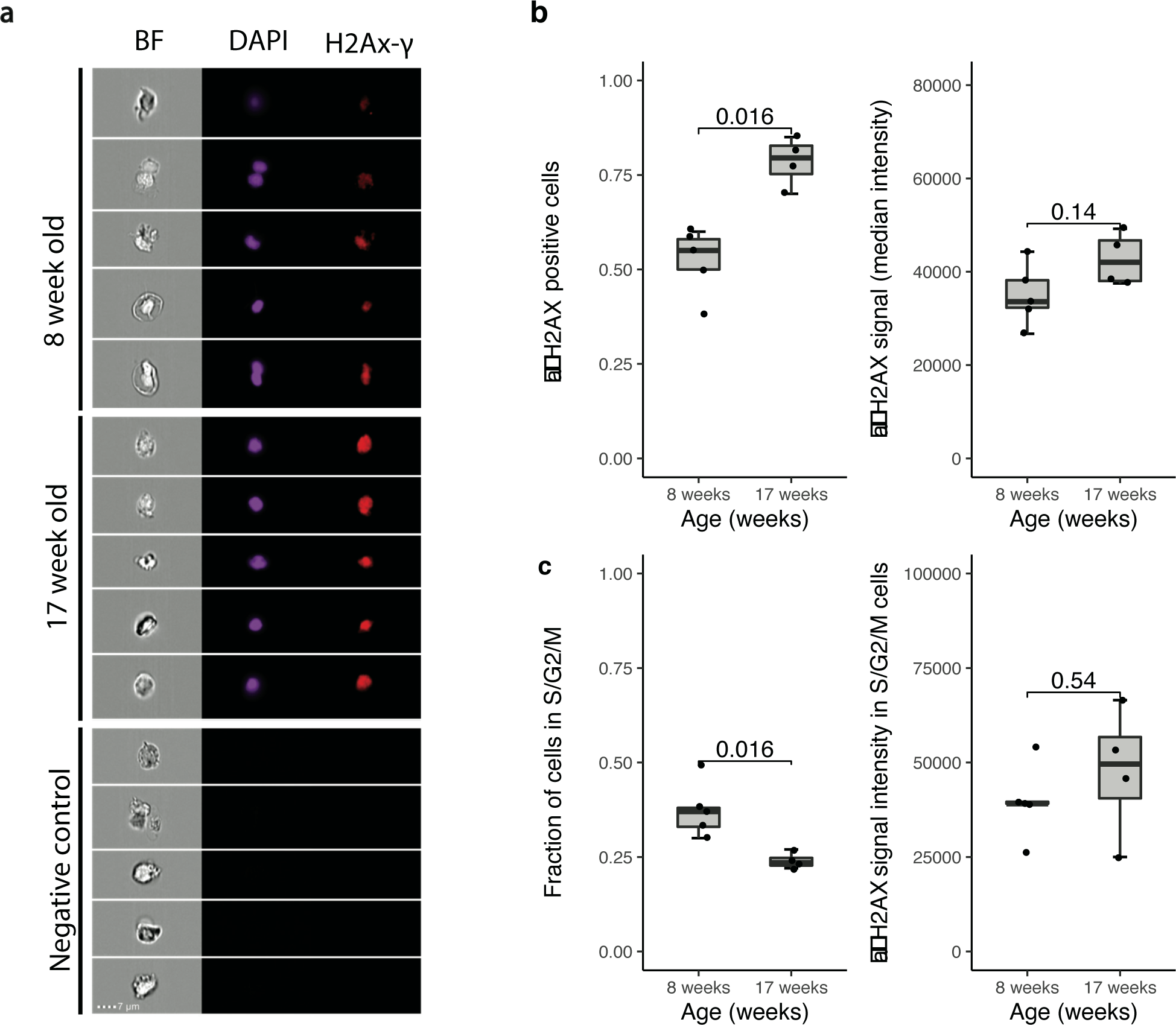
Proliferation impact on genomic stability within the turquoise killifish hematopoietic niche during aging: a) Immuno-fluorescence staining on single cell suspension isolated from the kidney marrows of adult (8-week-old) and aged-adult (17-week-old) killifish. Nuclei were stained with DAPI and anti-γ-H2AX antibody was used to mark double-strand breaks sites. All the data were recorded using Amnis ImageStreamX MkII Imaging Flow Cytometer and analysed in Amnis IDEAS v.6. b) Boxplots showing fraction of γ-H2AX positive cell and their median signal intensities among them for immune cells isolated from the kidney marrow of 5 adult and 4 aged-adult fish. c) Boxplots showing fraction of proliferating cells in S/G2/M phase measured based on their DAPI content and γ-H2AX signal intensities among these cells in 5 adult and 4 aged-adult fish. All the statistics were done using non-parametric Wilcoxon ranked sum test (two sided).

